# Metagenomic Insights into the Spatio-temporal Variation of Metal and Antibiotic Resistance Genes in Lake Erie

**DOI:** 10.1101/2024.10.02.616392

**Authors:** Saahith Reddy, E. Anders Kiledal

**Author notes:** **Address Correspondence to:** Anders Kiledal1, 100 North University Avenue Ann Arbor, MI 48109-1005.

## Abstract

Antibiotic resistance and metal toxicity in freshwater bodies have human health impacts and carry economic implications worldwide. The presence of metal and antibiotic resistance genes within microbial communities can be informative about both issues. The Laurentian Great Lakes contain nearly 20% of the world’s supply of freshwater; however, it is unclear how these genes are changing over time in this system. In this study, we characterized these genes in nearly two hundred metagenomes collected from multiple sites in western Lake Erie at a five-year time interval: 2014-2019. 11 metal resistance genes (MRGs) and nine antibiotic resistance genes (ARGs) were characterized and demonstrated significant changes in diversity and spatial distribution. Increased abundance was observed for genes like *aac(3)* and *TEM-1B* conferring resistance to aminoglycoside (gentamicin) and β-lactam antibiotics, respectively. MRGs associated with mercury, lead, and arsenic also increased in abundance over the five years. Collectively, our data point to a notable increase in both ARGs and MRGs in Lake Erie over five years, with a specific and significant increase in the abundance of genes conferring resistance to aminoglycoside and β-lactam antibiotic resistance and mercury contamination. Future integrated and systematic freshwater microbiome and public health investigations are needed to assess the potential impact on humans and environmental health from increasing microbial antibiotic and metal resistance in large freshwater reservoirs like the Great Lakes.

**IMPORTANCE:** Antibiotic and metal resistance genes (ARGs and MRGs) in microbial communities of the Laurentian Great Lakes have significant human and environmental health implications. However, an assessment of the Great Lakes’ microbiome for ARGs and MRGs is lacking. The abundance of 11 MRGs and 9 ARGs was characterized between 2014 and 2019 and showed significant abundance differences. Specifically, we observed an increase in genes conferring resistance to aminoglycoside (gentamicin) and β-lactam antibiotics (amongst the most commonly utilized antibiotics in humans), such as *aac(3)* and *TEM-1B,* respectively. MRGs conferring resistance to mercury, lead, and arsenic also increased in abundance, with the largest increase observed for mercury resistance genes such as *MerA, MerP,* and *MerT*. Collectively, these findings point to a concerning increased abundance of both ARGs and MRGs in Lake Erie. Further studies to assess the causes for the increase and the direct impact on human and environmental health are needed.

## INTRODUCTION

Since the introduction of modern water treatment standards and methodologies like chlorination, the threat of water-borne infectious diseases has significantly decreased in countries with access to clean water (1, 2). However, industrialization and increased environmental pollution, alongside the discovery and widespread use—and overuse—of antibiotics (3), combined with ineffective economic and public policies, have led to new health hazards affecting populations worldwide (4). These hazards include ramifications from both non-infectious toxicities related to contamination with metals such as mercury, arsenic, lead, and also microbial infections in humans driven by increased antibiotic resistance through the transmission of ARGs (antibiotic resistance genes) which pose a major global public health challenge (5, 6). Furthermore, metal and antimicrobial resistance genes are often co-selected due to similar resistance mechanisms (7–9).

Since the discovery of penicillin, routine use of antibiotics, particularly in agriculture and livestock farming, has led to a proliferation of bacteria capable of resisting their effects via antimicrobial resistance genes (ARGs) which are frequently horizontally transferred (10–13). While antibiotic resistance is a natural phenomena employed by microbes, the increase in clinically relevant antimicrobial resistance (AMR) from exposure to antibiotics used in healthcare and agriculture pose a serious threat to public health global, as highlighted by the World Health Organization (WHO) which ranked antimicrobial resistance among the top ten global public health threats confronting humanity (4, 14). Furthermore, a RAND Corporation analysis from nearly a decade ago estimated that, without action against AMR the decrease in global working-age populations by 2050 could be between 11-444 million people (5). ARGs encode proteins that resist antibiotics via a variety of mechanisms such as efflux pumps, antimicrobial inactivation, modification of the antimicrobial target, or limitation of antimicrobial uptake (15) and are present in various bacteria, with horizontal gene transfer of these genes leading to an ever-expanding repertoire of resistant organisms (16). This genetic exchange of resistance genes occurs via mechanisms such as natural transformation, conjugation, or bacteriophage-mediated transduction (17, 18). The introduction and excessive use of antibiotics have selected for and spread ARG-containing microbes, which now permeate environmental microbial communities, extending beyond healthcare settings and into different environments including freshwater systems (4, 10). Though environmental reservoirs such as contaminated waterbodies are known to be important for the spread and proliferation of ARGs, it is unclear to what extent (19).

Heavy metals such as lead (Pb), arsenic (As), and mercury (Hg) can cause significant chronic and acute toxicity in children and in adults (20, 21). Metal resistance genes (MRGs), including those for As, Hg, and Pb) resistance, are prevalent in bacterial communities across diverse habitats (21, 22). The role of MRGs in freshwater lake microbiomes remains poorly characterized (23). These metal resistance operons, found on transposons, genomic DNA, integrons, and bacterial plasmids, often associate with bacteria carrying ARGs (7, 24, 25). Moreover, like ARGs, MRGs can move and transfer among bacterial communities through horizontal gene transfer (17). The evolution and spread of ARGs are influenced not only by antibiotics but also by other factors; because of overlap in metal and antibiotic resistance mechanisms—particularly efflux pumps (26)—selective pressure from exposure to heavy metals can co-select for ARGs and contribute to their spread (7–9). MRGs can exert widespread and persistent co-selection pressure for ARGs through genetic linkages (27). Metagenomic studies that investigate the co-occurrence and spread of ARGs and MRGs in freshwater microbiomes remain poorly explored.

The Laurentian Great Lakes, a major interconnected freshwater system spanning the US and Canada, hold nearly 20% of the world’s freshwater. This unique ecosystem is characterized by dynamic and varied microbial communities, variations in mineral toxicity, and complex interactions that influence both environmental health and human well-being (28–31). The Great Lakes have been impacted by mercury contamination, with an extensive monitoring effort identifying source distributions and generally decreasing trends, with localized increases (29). Herein we present a time-resolved analysis of microbial communities from Lake Erie focusing on ARGs and MRGs. Utilizing high-throughput metagenomic shotgun sequencing of the Lake Erie microbiome, we evaluate microbial ARGs and MRGs, and identify an increase in ARGs related to aminoglycoside and β-lactam antibiotic resistance and also an increased abundance of MRGs, especially mercury resistance genes over a period of 5-years from 2014-2019. Thus, our data provide new insights into how this critical water system may be impacted by the growing pandemic of AMR and the potential contribution of metal toxicity (4, 32).

## RESULTS

### Microbial community structure of Lake Erie

Metagenomic sequences from samples collected throughout the western basin of Lake Erie (Supplemental Table 1) in 2014 and 2019 were obtained from the Great Lakes Atlas of Multi-omics Research (GLAMR) database (greatlakesomics.org). Community composition for each of the samples was also obtained from GLAMR, and average community compositions for samples from the two years were compared (Figure 1). Samples from 2014 on average had a higher abundance of Eukaryotic and Archaeal taxa, Alphaproteobacteria, and Cyanobacteria such as *Microcsytis*, responsible for annual harmful algal blooms.

**Figure 1:**
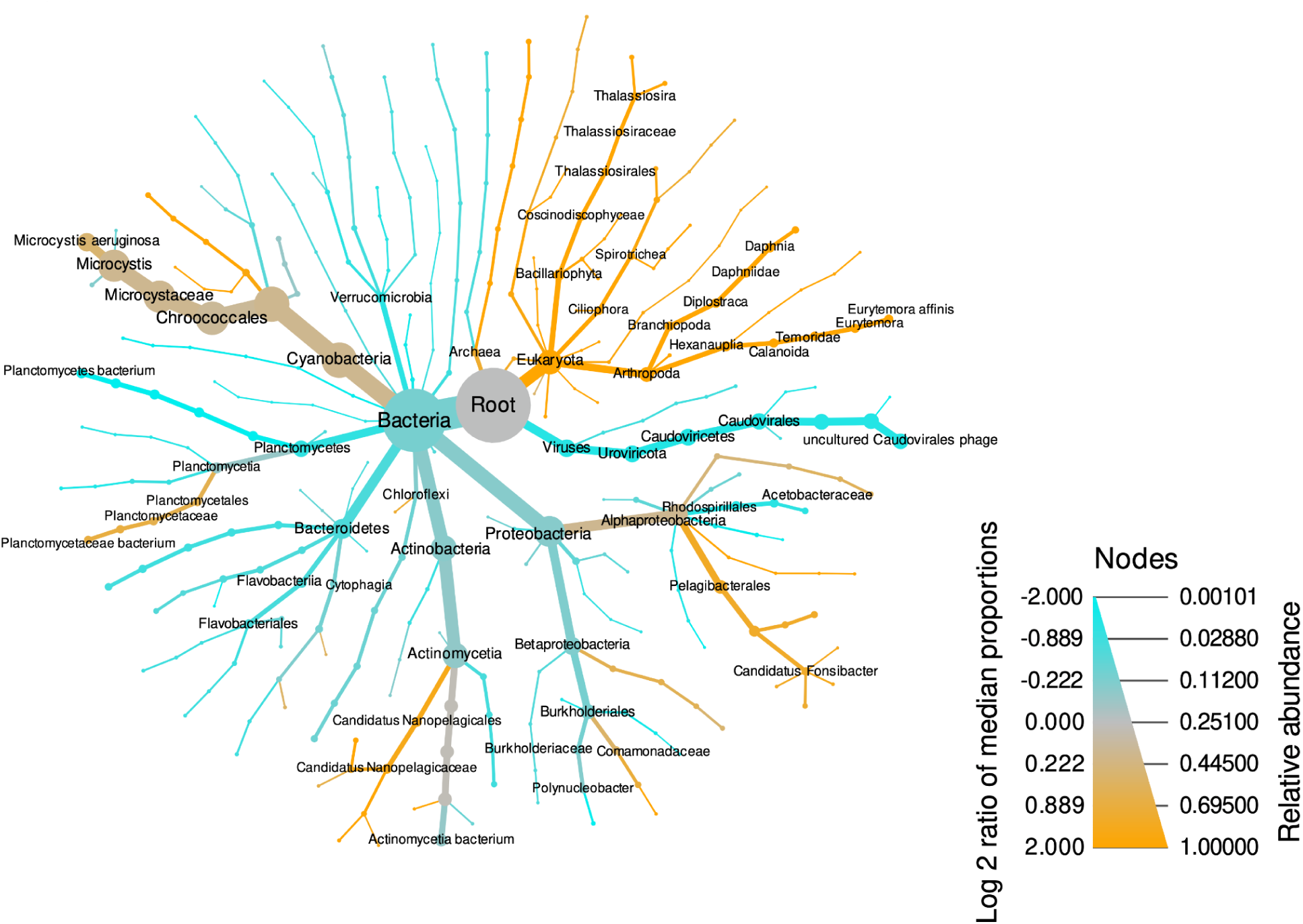
Community composition. Community composition was compared for metagenomes collected from western Lake Erie in 2014 and 2019. Composition was estimated based on normalized abundance (TPM) and taxonomic labels determined by LCA of hits to the UniRef100 database for contigs in metagenomic assemblies, obtained from the GLAMR database (greatlakesomics.org). Node size indicates the average abundance of a taxon, and node color indicates the log2 abundance ratio between the years, with taxa more abundant in 2014 shown in orange and those more abundant in 2019 shown in cyan.

### Antibiotic Resistance Gene Abundance

Metagenomic reads from each of the samples were mapped to reference antibiotic and metal resistance genes (ARGs and MRGs, respectively). Out of the 15 ARGs considered, 9 were identified in the Lake Erie microbiome (Figure 2A). Spatial differences were observed, although their significance is unclear due to the low number of observations. Genes conferring resistance to cephalosporin, fosmidomycin, gentamicin, glycopeptides, and piperacillin were widely distributed in the western basin of Lake Erie. Carbapenem ARGs were more abundant in samples collected closer to the Detroit River, whereas ARGs associated with resistance to ampicillin and aztreonam were generally found in samples closer to the shore and/or the Maumee River mouth.

**Figure 2.**
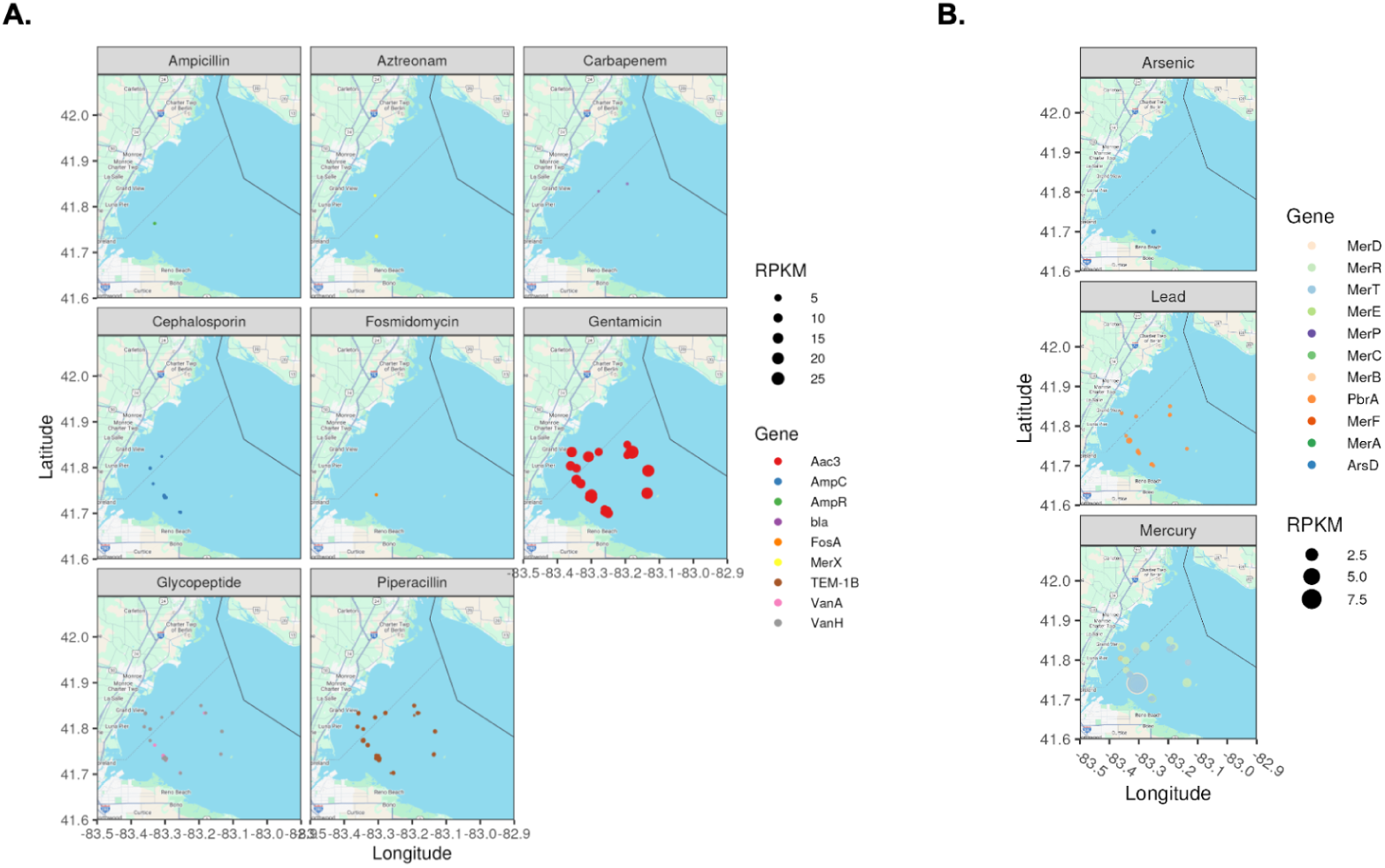
Distribution and abundance of ARGs and MRGs in western Lake Erie metagenomes. The abundance and spatial distribution of (A) antibiotic resistance genes (ARGs) and (B) metal resistance genes (MRGs) in western Lake Erie, determined by mapping metagenomic reads to reference gene sequences. Mapped read counts were normalized for sequencing effort and gene length. Panels correspond to genes grouped by the resistance conferred.

We next assessed whether this variation in the abundance of the seventeen ARGs changed over the 5 years from 2014 when compared to 2019. The relative abundance of the aminoglycoside (gentamicin) resistance gene, *Aac(3)* is notable as it was the most abundant of the ARGs in both 2014 and 2019, although more abundant in 2019 (Figure 3A) and also found in a higher proportion of samples (Figure 3B). Several ARGs were observed in 2019 but not in 2014, including ARGs conferring resistance to ampicillin, aztreonam, carbapenem, fosmidomycin, and glycopeptides (Figure 3B). In addition, we observed changes in abundance of the individual ARGs in a given year, but the variation in the ARGs between 2014 and 2019 were distinct (Figure 3C). The overall abundance of ARGs was compared between the two years (Figure 3D), showing that the median ARG abundance was lower in 2019 (p=0.11, Wilcox test), however this was largely due to an increased diversity of less abundant ARGs, as each individual ARG analyzed was either more abundant or more prevalent in 2019 samples. Thus in the five years the overall abundance and diversity of ARGs increased, including genes conferring resistance to β-lactam antibiotics (ampicillin, cephalosporin, piperacillin, and carbapenem), aminoglycoside (gentamicin), glycopeptide (vancomycin), and others such as polymyxin and fosmidomycin.

**Figure 3.**
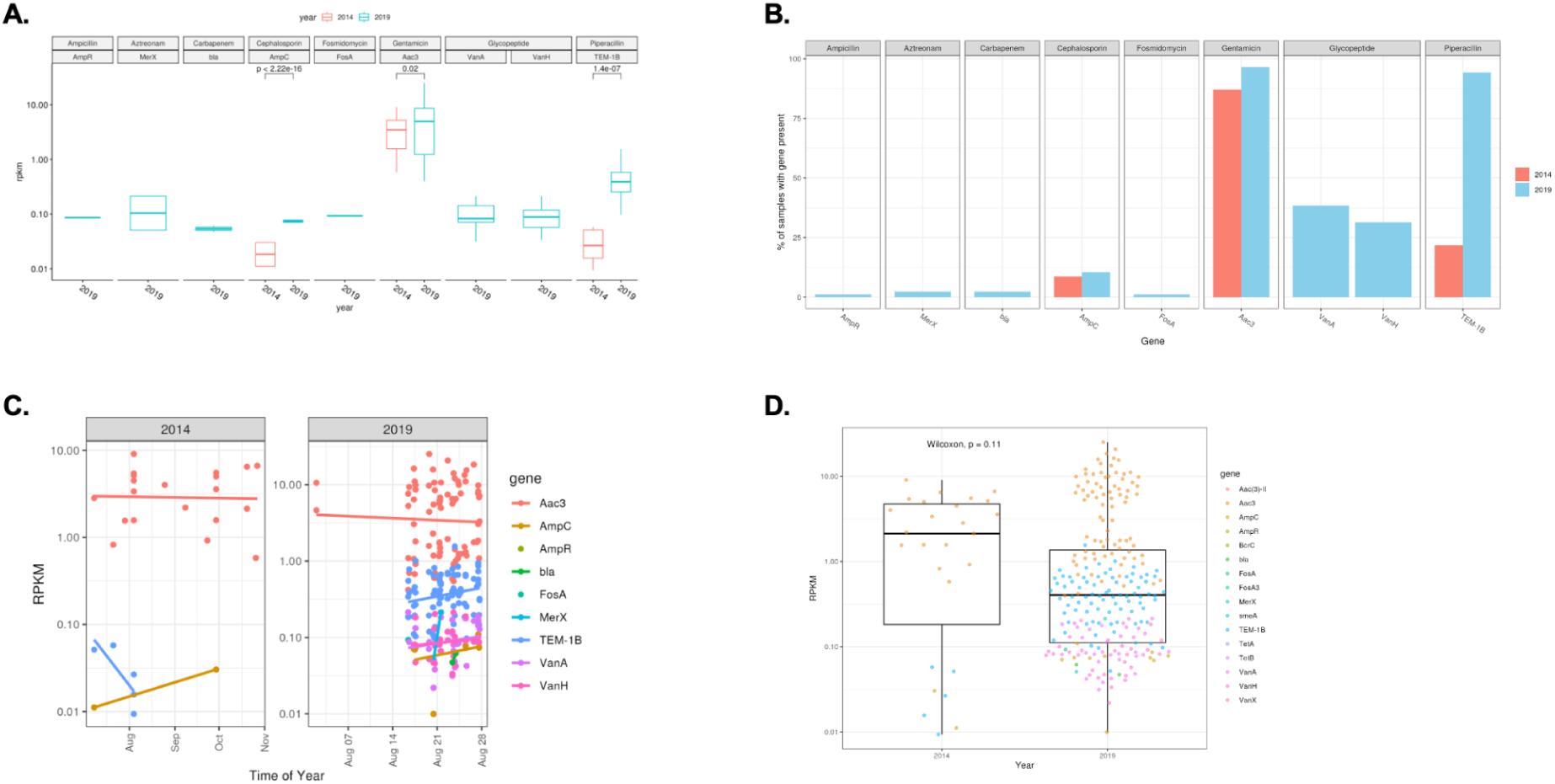
Abundance and prevalence of antibiotic resistance genes in western Lake Erie. Normalized antibiotic resistance gene abundances (RPKM) were compared between samples from 2014 and 2019 (A). Where enough observations were available, abundances were compared with Wilcoxon signed-rank tests and p.values are shown. Similarly, the % of samples containing a given gene (B) was compared in samples from 2014 and 2019 showing an increased prevalence for nearly all genes, with several resistance genes observed in 2019 but not 2014 samples. Intra-annual trends in gene abundance are shown in panel C; trends are illustrated with linear regressions for each gene. Overall, antibiotic resistance genes were significantly more abundant in 2019 than 2014 (D).

### Metal Resistance Genes for Hg, Pb, and As in the Lake Erie microbiome

All twelve MRGs (one for arsenic, two for lead, and nine for mercury) searched for were identified in the Lake Erie microbiome (Fig. 2B). The identified MRGs were widely distributed, with the exception of the arsenic resistance gene *arsD,* and genes conferring resistance to mercury and arsenic were generally more abundant in the southwest, closer to the Maumee River mouth (Fig. 2B). Mercury resistance genes were the most abundant MRGs, particularly merC, merD, merE, merF, merP, merR, and merT (Figure 4A). merA—an oxidoreductase key to mercury detoxification that reduces Hg^2+^ to elemental mercury—was more abundant in 2019 samples (p=1.3×10-7, Wilcox test), and merB, required for detoxification of particularly toxic methylmercury, was found at similar levels in both years (p=1, Wilcox test) (Figure 4A). Furthermore, while most of the samples collected in 2019 contained MRGs (albeit to a variable level), many of the 2014 samples did not demonstrate presence of MRGs suggesting a greater geographical distribution of MRGs in 2019 (Figure 4B). Interestingly, the lead resistance gene *pbrA* was observed in a greater proportion of samples from 2014 (Figure 4B), but at a higher median abundance in samples from 2019 (Figure 4A) and showed an increased trend in the individual years of both 2014 and 2019 (Figure 4C). We next assessed the trends in the overall abundance of all MRGs and also the individual 12 MRGs in the microbiome analyzed between the 5 year time span to determine the direction of change from 2014 to 2019. As shown in Figure 4D, the overall abundance of the MRGs increased from 2014-2019 (p = 3.8e-07), demonstrating an increase in overall MRGs over the half decade.

**Figure 4.**
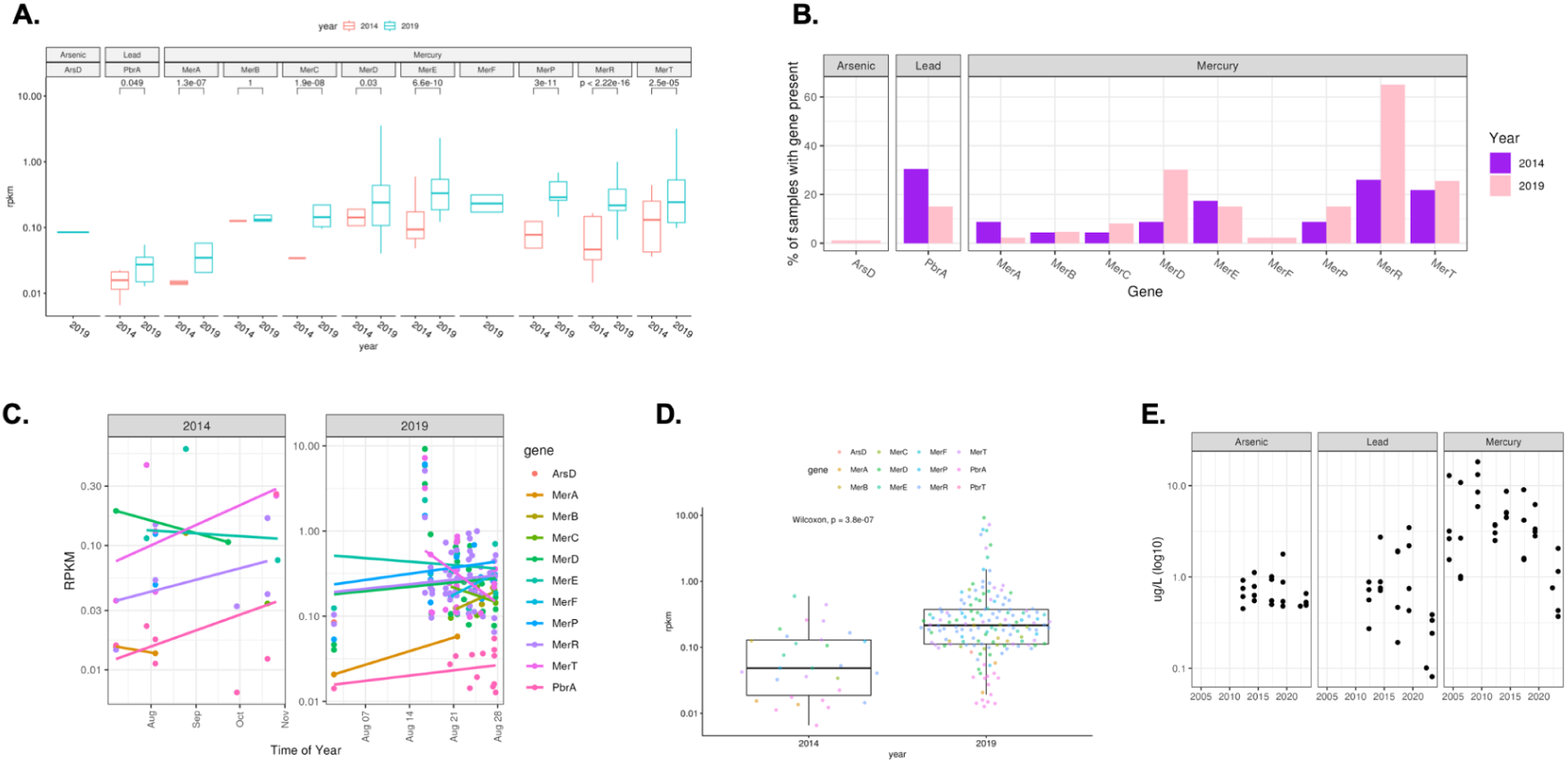
Abundance of metal resistance genes in western Lake Erie. The abundance of metal resistance genes (MRGs) was compared between samples from 2014 and 2019, revealing significantly more abundant lead and mercury resistance genes in 2019 samples (A). Where enough observations were available, abundances were compared with Wilcoxon signed-rank tests and p.values are shown. Similarly, the % of samples containing metal resistance genes were compared between samples collected in 2014 and 2019 (B). Intra-annual trends in metal resistance gene abundance are shown; linear regressions are shown for each gene (C). Overall, metal resistance genes were more abundant in 2019 samples than 2014 samples (D). For comparison, contemporaneous measurements of metal concentrations in western Lake Erie corresponding to the resistance genes analyzed are shown over the same time period (E) (DOI: 10.25976/6fgn-0915).

Contemporaneous measurements of arsenic, lead, and mercury concentrations were obtained from an Environment and Climate Change Canada (ECCC) monitoring dataset (33) to compare with resistance gene abundances (Figure 4E). Mercury concentrations were the highest, which matched gene results, followed by lead and arsenic. Lead resistance genes were generally observed at lower abundances than other MRGs, but were present in a larger proportion of samples potentially in keeping with environmental metal concentrations. Mercury and lead concentrations were variable, although they declined over the study period, while arsenic concentrations were nearly unchanged.

We also evaluated correlations in the abundance of MRGs and ARGs (Figure 5). Weak correlations were observed between the abundance of mercury resistance genes and genes conferring resistance to gylcopeptide antibiotics like vancomycin, which has high clinical relevance as a last-resort option for treatment of certain Gram positive bacterial infections. Several ARGs were also correlated, including those conferring resistance to macrolide-lincosamide-streptogramin (MLS), glycopeptide, piperacillin, and aztreonam (Figure 5A). Interestingly, there was a negative correlation in the abundance of mercury resistance genes and gentamicin resistance genes (Figure 5B).

**Figure 5.**
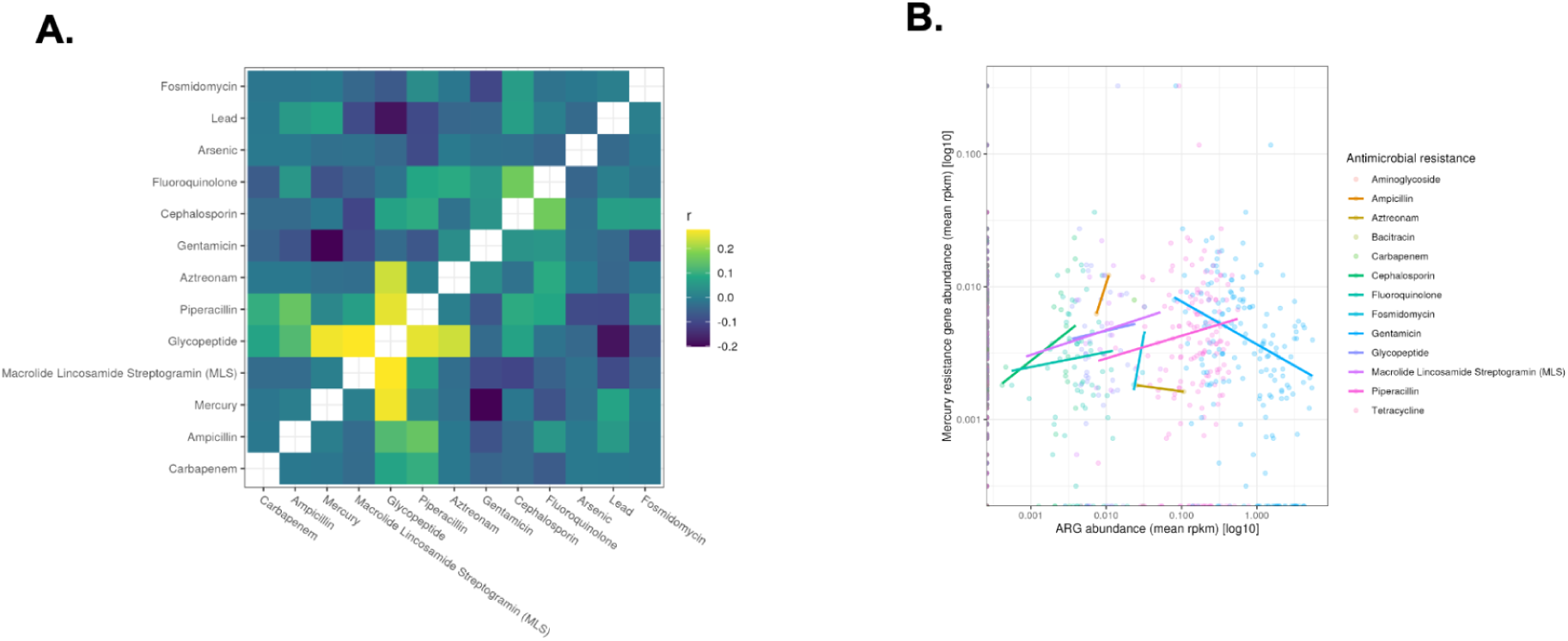
Correlation between metal resistance and antimicrobial resistance. The mean abundance of resistance genes were summarized by resistance target and pairwise Pearson correlations were calculated (A), showing a high degree of correlation between mercury resistance and resistance to a number of antimicrobials (B). However, the most abundant antimicrobial resistance genes (ARGs)–conferring resistance to gentamicin–were not correlated with mercury resistance, which had the most abundant metal resistance genes (MRGs.

## DISCUSSION

We investigated the microbial communities and the prevalence of ARGs and MRGs in the microbiome of Lake Erie at a five year interval, from 2014 and 2019. The sampling locations were similar to previous reports, thus providing for context to our observations and allowing them for future comparisons (34, 35). We observed spatial and temporal variations in microbial communities, ARGs, and MRGs, and identified nine ARGs and 11 MRGs with spatio-temporal fluctuations in the study period. Specifically, both ARGs and MRGs showed an overall increasing trend in the five years (Figure 3 & 4). The aminoglycoside (gentamicin) resistance gene *Aac3* were abundant and found in most samples, and over time an increase in β-lactam resistance ARGs was observed (Figure 3). ARGs corresponding to antibiotics with a high clinical relevance such as vancomycin and carbapenem were also observed (Figure 3). Likewise, mercury resistance genes, *MerA-F* and *MerP,R,T*, showed an increasing trend during the study period (Figure 4).

Resistance genes in microbial communities carry the potential for transfer to pathogenic bacteria through horizontal gene transfer mechanisms and are a significant threat to public and environmental health (36, 37). However, it is important to recognize that many bacterial species developed the capacity to withstand antibiotics and metals well before the mass production of antibiotics or contamination from the industrial wastes and fertilizers. A key driver of resistance before the widespread use of antibiotics or metals likely stems from the perpetual competition for natural resources among microorganisms (38, 39). This includes the natural production of secondary metabolites between the bacteria (38); notably gentamicin, which had the most abundant resistance genes, is produced by species in the Actinobacterial genus *Micromonospora* commonly found in freshwater systems (40). Additionally contamination of groundwater and soil through ancient geological processes also lead to development of MRGs as a consequence of natural geological processes (39, 41). But the advent of clinical development and overuse of antibiotics, fertilizers and industrial wastes have dramatically altered this landscape for resistance by introducing significant selective pressures, not only affecting human and domestic animal microbiota but also environmental microbiomes polluted with antibiotics, such as the Great Lakes (4, 5).

The ARGs and MRGs linkage is an increasingly appreciated issue in microbial communities (42). Mobile genetic elements like plasmids and transposons often harbor both metal and antibiotic resistance genes, suggesting that a single selective pressure can simultaneously enrich for both traits (42). Interestingly, our findings show a general increase in ARGs and MRGs trends over the past decade, especially for the aminoglycoside-resistant gene *Aac3* and beta-lactam resistance gene TEM-1B, and the mercury resistance genes *MerB, MerD, MerT and MerR*. The observed trend in MRGs is consistent with some findings in the levels of mercury in fish (43–46), although mercury concentrations are generally thought to be declining in the Great Lakes (29). Nevertheless, internal lake cycling or local hotspots could differ from general trends. Furthermore, while antibiotic resistance genes can be natural to microbes, the presence of certain ARGs, such as *Aac3-II* has been directly linked to hospitals with excessive use of aminoglycosides suggesting a potential reason for increase in ARG *Aac3* and *Aac3-II* in Lake Erie could be contamination from clinical use of antibiotics (47). These data suggest that the Lake Erie ecosystem might have developed increased concurrent selective pressures on aminoglycoside, beta-lactam and mercury resistance and may have implications for the simultaneous spread of mercury, beta-lactam and aminoglycoside resistance. Given the major role played by plasmids in propagation of gram negative bacterial resistance genes (48), our data raises the hypothesis that the mobile genetic elements for certain ARGs such as *Aac3* and the MRGs, such as *MerD/A* genes may be similar. However, we only observed weak correlations between the abundance of MRG and ARG abundance (Figure 5).

Furthermore, high levels of metals in urban areas can amplify the co-selection of MRGs and ARGs, while the concurrent transfer of resistance genes through MGEs can facilitate the persistence and dissemination of ARGs (49–51). The patterns of ARGs and MRGs in urban lakes are associated with prolonged human activities and eutrophic conditions of the lakes (52). Microbial metal resistance mechanisms can be indicators of metal contamination (i.e. biosensors) (53, 54), and so an increasing trend might be indicative of metal contamination in the lake. Metals have long been known to possess antibacterial properties (‘metalloantibiotics’) (55) and evidence indicates that silver (Ag) and mercury (Hg) are highly toxic to most bacteria and exhibit microbicidal activity even at very low concentrations (56). MRGs can also be used in the development of bioremediation and environmental engineering strategies that involve modifying the microbiome (57). Specifically, the taxonomic and functional microbiome datasets from our study could provide essential information for assessing the feasibility of bacteria-mediated bioremediation of various metals and minerals. It may help in potential development of engineering processes that leverage multi-metal-resistant bacteria to diminish metal toxicity in the environment (57, 58).

Our study, however, has limitations and requires correlation with epidemiological and clinical studies to better quantify effects on human health. Moreover, our metagenomic data does not provide information about the expression of resistance genes which would provide more specific insights into conditions and selective pressures experienced by resident microbes. Furthermore, we considered data from two years which limits inference about long-term trends as the years included might not be representative of longer-term trends (or lack of trends). Our study also did not investigate the specific drivers for the type or degree of ARG and MRG changes in Lake Erie. The observed results could be a reflection of a combination of reasons such as environmental protection laws, waste handling practices, industrial pollution and widespread use of antibiotics. Future studies will need to determine the general and the specific causes for the general increasing trends of ARGs and MRGs, especially the aminoglycoside and the β-lactam resistance genes and the mercury resistance genes (59–61). Despite these limitations, our findings provide insights with potential implications for Great Lakes environmental policy; regulations may need to extend beyond the mere presence of chemical pollutants, considering their potential to select for resistance traits against heavy metals and antibiotics in microbial communities. The CDC tackles antimicrobial resistance through its Antimicrobial Resistance Solutions Initiative, which employs a One Health method to enhance healthcare, public health, veterinary medicine, agriculture, food safety, as well as research and manufacturing (62, 63). Our study underscores the importance of maintaining Lake Erie’s ecological integrity to safeguard public health and highlights the potential for co-selection and the spread of metal and antibiotic resistance through freshwater lakes.

## MATERIALS AND METHODS

### Identification of metagenomic samples

Publicly available shotgun metagenome samples from the Western Basin of Lake Erie were identified using the Great Lakes Atlas of Multi-omics Research (GLAMR) (greatlakesomics.org). Associated metadata for these samples was also collected and used for subsequent analysis (Supplemental Table 1). Sample collection information is described in manuscripts associated with original data publication (35, 64, 65). Methods for measurement of environmental conditions and nutrient concentrations associated with a subset of the samples collected by the NOAA Great Lakes Environmental Research Laboratory (NOAA-GLERL) and the Cooperative Institute for Great Lakes Research (CIGLR) have also been previously described (34, 35).

### Bioinformatics processing of metagenomic samples

Initial processing of metagenomic samples was conducted using the GLAMR bioinformatics pipelines (github.com/Geo-omics/GLAMR_omics_pipelines). Raw sequence reads were quality filtered, deduplicated, and quality and adapter trimmed using fastp v0.23.2 (parameters: --dup_calc_accuracy 6 --detect_adapter_for_pe --cut_front --cut_tail --cut_window_size=4 --cut_mean_quality 20 --length_required 50 --n_base_limit 5 --low_complexity_filter --complexity_threshold 7) (66). BBmap was used to remove human contaminant reads that mapped to the GENCODE release 38 human genome (67).

### Community composition summary

Taxonomic abundance information was obtained from the GLAMR database, calculated by summarizing contig abundance (TPM, calculated with the coverM tool and minimap aligner (68, 69)) and taxonomy determined with mmseqs2 and the UniRef100 database (70, 71). Figures were produced with the metacodeR R package (72).

### Quantification of antibiotic and metal resistance genes

Antibiotic and metal resistance genes were identified in several publications and representative sequences were obtained from GenBank (Supplemental Table 2) (73). Quality controlled metagenomic reads were mapped to these nucleotide sequences using MiniMap2 (69), filtered using filterBam from Augustus (minCover = 50, minID = 80), and processed with Samtools (74). Counts of mapped reads were determined using the Rsamtools package version 2.20.0 (75) and R version 2.1.2 (76). Custom R scripts were used to calculate normalized abundance values (RPKM) based on the total number of sequenced reads per sample, the reference gene length (kb), and the number of mapped reads that passed filtering criteria.

### Data summarization, statistics, and figure generation

Data was summarized using custom R scripts, leveraging packages in the tidyverse where appropriate (76, 77). The ggpubr R package version 0.6.0 was used to run non-parametric Wilcox and Kruskal Wallis tests and generate figures. Other figures were produced using the ggplot2 R package version 3.5.1 (78).

### Data and code availability

Shotgun metagenomic data was obtained from the GLAMR database, which in turn obtains most sequences from the National Center for Biotechnology Information (NCBI) Sequence Read Archive (SRA) database. Sample identifiers are included in Supplemental Table 1. Code used to conduct the analyses is available on GitHub (github.com/akiledal/Erie_MRG_ARG).

## ACKNOWLEDGMENTS

This work was supported by funding from the NOAA Great Lakes Omics program distributed through the UM Cooperative Institute for Great Lakes Research (NA17OAR4320152 & NA22OAR4320150).

No regulatory approvals were required.

## Supplemental items

**Supplemental table 1: Metagenomic samples and metadata used for analyses.**

Shotgun metagenome samples from the Western Basin of Lake Erie were identified using the Great Lakes Atlas of Multi-omics Research (GLAMR) (greatlakesomics.org). Associated metadata for these samples is also included. Includes latitude and longitude (decimal degree) coordinates for each station, depth, temperature, and several other values.

**Supplemental table 2: Description of metal and antibiotic resistance genes analyzed.**

All gene information was obtained from GenBank.

